# Faecal microbiota transplant from aged donor mice affects spatial learning and memory via modulating hippocampal synaptic plasticity- and neurotransmission-related proteins in young recipients

**DOI:** 10.1101/866459

**Authors:** Alfonsina D’Amato, Lorenzo Di Cesare-Mannelli, Elena Lucarini, Angela L. Man, Gwenaelle Le Gall, Jacopo J. V. Branca, Carla Ghelardini, Amedeo Amedei, Eugenio Bertelli, Mari Regoli, Alessandra Pacini, Giulia Luciani, Pasquale Gallina, Annalisa Altera, Arjan Narbad, Massimo Gulisano, Lesley Hoyles, David Vauzour, Claudio Nicoletti

## Abstract

**Background:** The gut-brain axis and the intestinal microbiota are emerging as key players in health and disease. Shifts in intestinal microbiota composition affect a variety of systems, however, evidence of their direct impact on cognitive functions is still lacking. We tested whether faecal microbiota transplant (FMT) from aged donor mice into young adult recipients affected the hippocampus, an area of the central nervous system (CNS) known to be affected by the ageing process, and related functions.

**Methods and Findings:** Young adult mice were transplanted with the microbiota from either aged or age-matched donor mice. Following transplantation, characterization of the microbiotas and metabolomics profiles along with a battery of cognitive and behavioural tests were performed. Label-free quantitative proteomics was employed to monitor protein expression in the hippocampus of the recipients. Gut permeability, levels of circulating cytokines and expression of markers of microglia cells were also assessed. FMT from aged donors led to impaired spatial learning and memory in young adult recipients, whereas anxiety, explorative behaviour and locomotor activity remained unaffected. This was paralleled by altered expression of proteins involved in synaptic plasticity and neurotransmission in the hippocampus. Also, a strong reduction of bacteria associated with short-chain fatty acids (SCFAs) production (*Lachnospiraceae, Faecalibaculum*, and *Ruminococcaceae*) and disorders of the CNS (*Prevotellacea*e and *Ruminococcacea*e) was observed. Finally, microglia cells of the hippocampus fimbria, acquired an ageing-like phenotype, while gut permeability and levels of circulating cytokines remained unaffected.

**Conclusions:** These results demonstrate a direct effect of the age-associated shifts of the microbiota on protein expression and key functions of the central nervous system. Furthermore, these results additionally highlight the paramount importance of the gut-brain axis in ageing and provide a strong rationale to devise therapies aiming to restore a young-like microbiota to improve cognitive functions in the elderly.

## Introduction

Ageing is an inevitable process that starts immediately after birth and ultimately leads to the loss of functional capacity in several body systems, including the cardiovascular, skeletomuscular, osteoarticular and neuro-immune-endocrine, and is often associated with a decline in psychological wellbeing and cognitive function. In the past few years it has been brought to the surface that events taking place in the gut play an important role in the ageing process [1] and recently, the existence of bidirectional communication between the gut and the brain – the gut-brain axis – has emerged as an important player in shaping aspects of behaviour and cognitive function [2]. In particular, the gut microbiome has been reported to play an important role within this scenario. A modest alteration in the composition of the gut microbiota, induced by either diet or antibiotics, is sufficient to cause changes in mouse brain chemistry and function [3, 4]. In particular, the oral administration of antibiotics resulted in an increase in brain-derived neurotrophic factor (BDNF) in the hippocampus. Furthermore, reduced expression of the synaptic plasticity-related genes PSD-95 and synaptophysin, important in memory formation and maintenance, were also reported [5]. Yet, germ-free (GF) mice are less responsive to exposure of environmental stressors compared to conventionally reared mice, with conventionalisation of GF animals impacting significantly on brain development [6]. This observation, along with a growing body of evidence, has shown that the gut microbiota plays a major role in in the development and function of the CNS, affecting learning and memory via metabolic, neuroendocrine and immune pathways [7]. In addition, dysbiosis has been associated with a variety of neurological disturbances ranging from depression [8] to autism [9, 10], along with neurodegenerative diseases such as Parkinson’s disease and multiple sclerosis [11, 12]. Not surprisingly, faecal microbiota transplant (FMT) is being investigated as a therapeutic option not only for GI-tract-related diseases [13], but also for CNS disorders [14, 15]. Thus, it is plausible to hypothesize that age-associated changes in the microbiota composition have a direct effect on the CNS, potentially contributing to the decline of cognitive function seen in the elderly. Recently, it has been reported that FMT from aged donor mice into young germ-free (GF) recipients increased hippocampal neurogenesis, intestinal growth, and activation of the prolongevity signaling pathways in the liver [16]; however, up to date the direct impact of the ageing microbiota on cognitive functioning and activity has not been addressed. We report that FMT from aged mice led to a decline of spatial learning and memory in young adult (henceforth termed adult) recipients via modulation of synaptic plasticity- and neurotransmission-related proteins in the hippocampus, an area of the CNS known to be heavily affected by the ageing process [17]. The impact of age-associated shifts of microbiota on the CNS was further confirmed by the observation that microglia cells of the hippocampus fimbria of adult mice acquired an ageing-like phenotype after FMT from aged donors. These changes occurred in the absence of an overt increase of both plasma levels of inflammatory cytokines and intestinal permeability.

## Materials and methods

### Animals and brain sample preparation

Adult (3 month) and aged (24 month) male C57BL/6 mice were used. Experiments were conducted under the guidelines of the Scientific Procedure Animal Act (1986) of the U.K. and approved by the Ethical Review Committee of the University of East Anglia, Norwich (AWERB ref: 70/7583) or at the Ce.S.A.L (Centro Stabulazione Animali da Laboratorio, University of Florence, Italy) under the approval of the Italian Ministry of Health (No. 54/2014-B) and the Animal Subjects Review Board of the University of Florence. Mice (Envigo, Varese, Italy or Charles River, UK) were provided with food and water ad libitum. Environmental temperature was kept at 23±1 °C with a 12 h light/dark cycle. All efforts were made to minimize animal suffering and to reduce the number of mice used.

### Faecal material preparation and FMT regime

Faecal material was collected from adult and aged mice and placed into Eppendorf tubes containing 500 ml of freezing solution (sterile saline solution with 12.5% glycerol), and homogenized. The suspended pellets were then stored at −80°C until utilized. For FMT, mice were randomized into the following groups (n=12 per group): control (no antibiotic treatment, no FMT); FMT-adult (antibiotic treatment followed by FMT from adult age-matched donors), FMT-aged (antibiotic treatments followed by FMT from aged donors). Antibiotic mix was administered by oral gavage. The antibiotic/anti-fungal regime was as follows: day 1-3 mice were gavaged daily with anti-fungal treatment with amphotericin B 1 mg kg-1, day 4-17 mice received a daily gavage of metronidazole 100 mg kg^-1^ while the antibiotic mix (ampicillin 1 g L^-1^, vancomycin 0.5 g L^-1^ and neomicin 1 g L^-1^ was added to drinking water), day 18-24 daily oral gavage with ampicillin 1 g L^-1^, vancomycin 0.5 g L^-1^, neomicin 1 g L^-1^, metronidazole 100 mg kg^-1^, amphotericin B mg kg^-1^. FMT was carried out via oral gavage with a faecal suspension (100 mg ml^-1^) in a final volume of 150 ml. FMT was performed six times on days 24-28 and 35 from the beginning of antibiotic regime.

### DNA extraction, amplicon sequencing and analyses of 16S rRNA gene sequence data and metabolomics analysis

Pre- and post-FMT faecal material was used for microbiota profiling. DNA was extracted from faecal samples using the FastDNA SPIN Kit for Soil (MP Biomedicals) with three bead-beating periods of 1 min as previously described [18]. DNA concentration was normalised to 1 ng/μL by dilution with DNA elution solution (MP Biomedicals, UK) to produce a final volume of 20 μL. Normalised DNA samples were sent to the Earlham Institute (Norwich, UK) for PCR amplification of 16S rRNA genes and paired-end Illumina sequencing (2 × 250 bp) on the MiSeq platform. The V4 hypervariable region of the 16S rRNA genes was amplified using the 515F and 806R primers with built-in degeneracy as previously reported [19, 20]. Sequence data were provided in fastq format. All processing and analyses were done in R/Bioconductor making use of the following packages: GEOquery 2.50.0 [21]; dada2 1.10.0 [22]; phyloseq 1.26.0 [23]; tidyverse 1.2.1 (https://www.tidyverse.org); vegan 2.5.3; viridis 0.5.1; msa 1.14.0 [24]; phangorn 2.4.0; ALDEx2 1.14.0 53; gplots 3.0.1. Taxonomy was assigned to chimera-free Exact Sequence Variants [25] using Silva 132 (downloaded from https://zenodo.org/record/1172783#.W-B0iS2cZBw on 5 November 2018). Data were filtered to remove undefined phyla and taxa present in fewer than two animals. Significance of differences between different diversity measures was determined using Wilcoxon rank sum test. ALDEx2 was used to determine statistically significant differences (Welch’s t test, Wilcoxon) between mouse groups. The 16S rRNA gene sequence data have been deposited in the NCBI BioProject database (https://www.ncbi.nlm.nih.gov/bioproject/) under accession number PRJNA524024.

Metabolites were analysed and quantified by ^1^H-NMR analysis as previously described [26]. Briefly, 20 mg of frozen faeces were thoroughly mixed on a vortex with 1 mL of saline phosphate buffer followed by centrifugation (18,000 g, 1 min). High-resolution 1H-NMR spectra were recorded on a 600 MHz Bruker Avance spectrometer fitted with a 5 mm TCI proton-optimized triple resonance NMR ‘inverse’ cryoprobe and a 60 slot autosampler (Bruker, Rheinstetten, Germany). Metabolites were identified using information found in the literature or on the Human Metabolome Database, http://www.hmdb.ca/ and by use of the 2D-NMR methods, (e.g COSY, HSQC and HMBC [27] and quantified using the software Chenomx® NMR Suite 7.0TM.

### Barnes Maze test

This test was conducted according to previously described method [28] All sessions were performed on a wet platform under a room lightning of 400 lux to increase the mouse aversion for the platform. Sessions were recorded using a video-tracking system (ANY-maze, Ugo Basile, Varese, Italy). The platform and the escape box were cleaned thoroughly between each mouse session and the surface of the platform was moistened again to avoid mice using olfactory cues to solve the task. Before performing the test the mice were trained to find the cage for 4 days with 2 sessions per day.

### Novel object recognition test

The novel object recognition task chamber was carried out as previously described [29]. For each mouse, the time spent interacting with each object during the acquisition and recognition phases was video recorded and analysed blind. The corrected mnesic index (or discrimination index) was calculated as follows: (time spent exploring the novel object - time spent exploring the well-known object)/total time spent exploring both the objects.

### Elevated plus maze

To evaluate anxiety-related behaviour, the elevated plus maze test was conducted as previously described [30]. Time spent in each kind of arms (sec) was recorded for 5 min. Percentage of entries into the open arms and that of time spent in the open arms were determined. Data acquisition and analysis were performed automatically using ANY-maze software.

### Open field test (OFT)

The OFT was conducted as previously described [31]. Briefly, mice were placed in the centre of the OFT and the total distance the mice travelled along with the time they spent in the centre of the field within 5 min was recorded with a video tracking system. The open field maze was cleaned between each mouse with 20% ethanol to eliminate odour.

### Liquid Chromatography-MS analysis

Each sample was analysed by nLC MS/MS Orbitrap Fusion trihybrid mass spectrometer coupled with a nano flow UHPLC system (Thermo Fischer Scientific, USA), via a nano electrospray source with an ID 0.01mm fused silica PicoTip emitter (New Objective). The peptides were separated, after trapped on a C18 pre-column, using a gradient of 3-40% acetonitrile in 0.1% formic acid, over 50 min at flow rate of 300 nL/min, at 40°C. The MS method consisted of a full scan in Orbitrap analyser (120000 resolution), followed by the combination of CID and HCD collisions. The peptides were fragmented in the linear ion trap by a data-dependent acquisition method, selecting the 40 most intense ions. Dynamic exclusion of sequenced peptides was set to 30s. Data were acquired using Xcalibur software (Thermo Scientific, USA). All analyses were performed in triplicate. MS raw data were analysed by MaxQuant (version 1.6.2.3), using Andromeda search engine in MaxQuant and consulting the Homo Sapiens UniProtKB database; the tolerance on parents was 10 ppm and on fragments was 0.02 ppm. The variable modifications allowed were oxidation on methionine and acetylation on N terminus and carbamidomethylation on cysteine as fixed modification. The false discovery rate was below 1%, using a decoy and reverse database, and a minimum number of seven amino acids were required for peptide identification. Proteins and protein isoforms were grouped into protein groups. Label-free quantitative analyses were also performed by MaxQuant software, using the MaxLFQ algorithm. The quantitation values were obtained on high-resolution three-dimensional peptide profiles in mass-to-charge, retention time and intensity space. The mass spectrometry proteomics data have been deposited to the ProteomeXchange Consortium via the PRIDE [32] partner repository with the dataset identifier PXD016432.

### Intestinal permeability and plasma cytokines

Plasma levels of cytokines were assessed by Enzyme-linked immunosorbent assays (ELISA) Assays were carried out according to the manufacturer’s instructions (all kits from eBioscience). Intestinal permeability was assessed using a method described previously [33]. Animals were orally delivered with 0.5 ml of PBS containing 25 mg of FITC-labelled dextran (FD4; Sigma-Aldrich) and blood samples were collected after 45 minutes. Plasma concentration of FD4 concentration was assessed using fluorescence spectrometer (LS 55 conducted with FL WinLab software, PerkinElmer, USA) at an excitation wavelength of 490 nm and emission wavelength of 520 nm.

### Statistical analyses

Microbiomic data (genus level) were correlated (Pearson) with metabolomic data using aldex.corr() within the Bioconductor package ALDEx2 [18]. For the MS study all data were evaluated by Perseus statistical software. The protein expression fold change variation between two groups was analysed by two sides t test, setting FDR less than 0.015 and s0 of 0.1. The differentially expressed proteins, with a significant ratio FMT-aged/FMT-adult (P value < 0.05), were analysed by Ingenuity Pathway Analyses (Qiagen) and only the differentially regulated pathway with a significant z score, after Bonferroni test, were considered. The symbols shown in the network are explained at http://www.qiagenbioinformatics.com/products/ingenuity-pathway-analysis. Behavioural measurements were performed for each treatment in two different experiments (n=10-12 mice/group). All assessments were performed in blind of the treatment received by the mouse groups. Results were expressed as means ± S.E.M. and the analysis of variance was performed by two-way ANOVA. A Bonferroni’s significant difference procedure was used as post-hoc comparison. P values of less than 0.05 or 0.01 were considered significant. Data were analyzed using the “Origin 9” software (OriginLab, Northampton, USA).

## Results

### Microbiota of adult recipients acquires an aged phenotype following FMT from aged donors

We investigated in the first instance the faecal microbiotas of adult (3 months) and aged mice (24 months) (Fig. 1) to select the donors for the FMT procedure. Measures of alpha diversity showed only a significant (P=0.0107) difference in species richness between the two cohorts, with the faecal microbiota of aged mice harbouring more different amplicon sequence variants (ASVs) than adult mice (Fig. 1a). Beta diversity (Bray Curtis) analysis showed clear separation of the two groups of mice based on their faecal microbiotas (Fig. 1b). Comparison of relative abundances of different taxa showed significantly more ASVs associated with *Ruminiclostridium, Butyricicoccus, Lachnoclostridium, Lachnospiraceae* spp., *Shuttleworthia* and *Marvinbryantia*, and significantly fewer associated with *Staphylococcus, Jeotgalicoccus, Facklamia, Parvibacter, Enterorhabdus, Muribaculum, Parabacteroides* and *Anaerostipes* in the adult mice relative to the aged mice (Fig. 1c). Integration of faecal microbiomic and metabolomic data showed an association between lower levels of faecal short-chain fatty acids (SCFAs) and decreased representation of obligate anaerobes such as the *Lachnospiraceae* and *Ruminococcaceae* (Supplementary Fig. S1 and S2).

**Figure 1.**
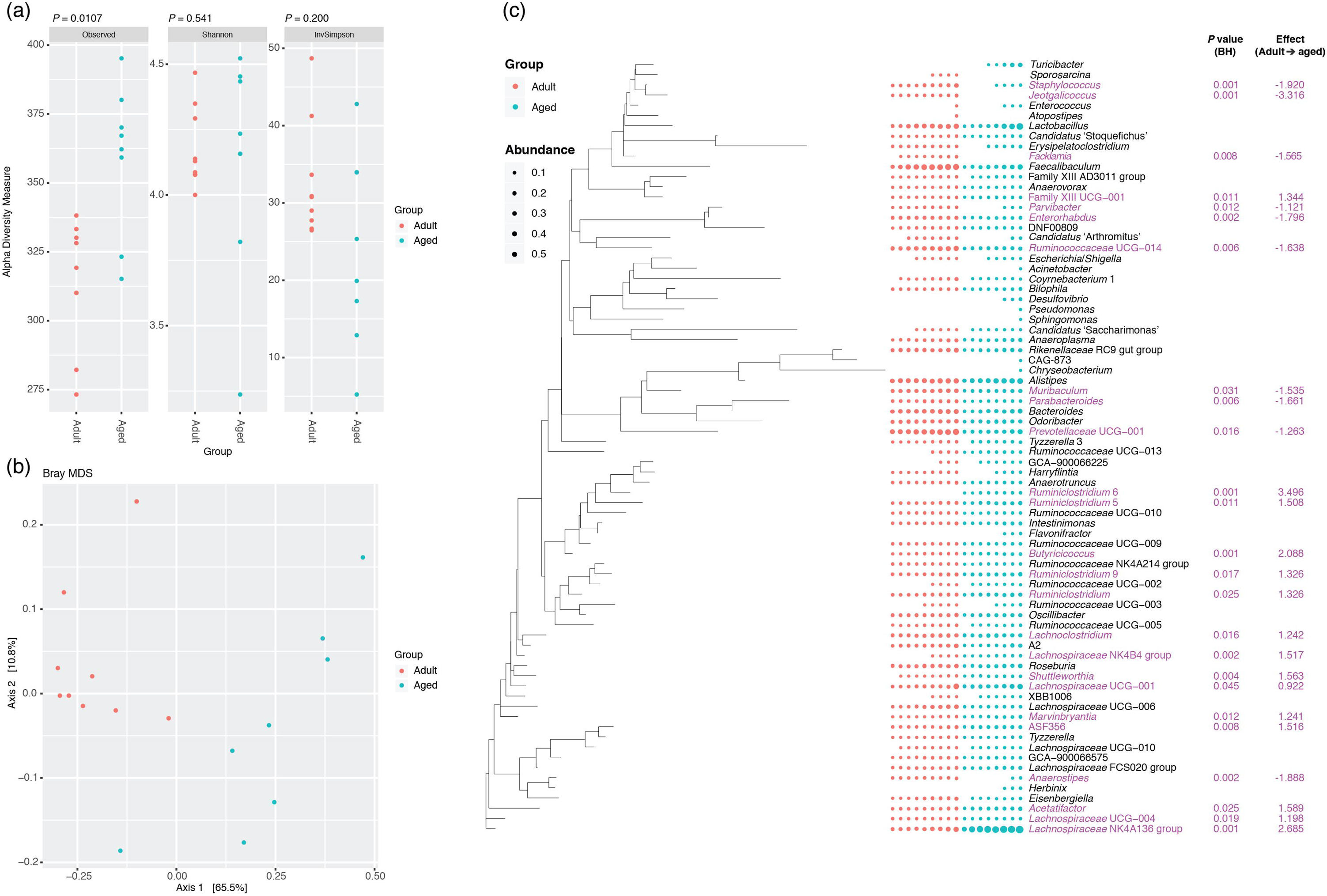
Comparison of the faecal microbiotas of adult and aged mice. (a) Measures of alpha diversity. Significance of differences between the two groups was assessed by Wilcoxon rank sum test. (b) MDS plot of a Bray Curtis assessment of beta diversity. Data presented are for ASVs present in more than two animals and prevalence threshold of 1 % at the genus level. (c) Comparison of the relative abundance of different genera present in the faecal microbiota of the two cohorts. Purple text, significantly different (Welch’s t test and Wilcoxon; P<0.05, Benjamini–Hochberg) based on ALDEx2 analyses.

Based on this analysis, faeces from three of the adults and two of the aged mice included in the analysis above were pooled and used as donors for FMT (Fig. 2). Adult mice received FMT either from adult donors (FMT-adult) or aged donors (FMT-aged). Measures of alpha diversity showed no significant differences between the two mouse sub-groups (Fig. 2a). Bray Curtis analysis showed clear separation of the pre- and post-FMT groups, with the post-FMT samples clustering with their respective inoculants (Fig. 2b). It is notable that antibiotic treatment of pre-FMT mouse 1 (Ms1) and mouse 2 (Ms2) was not completely successful as they clustered more closely with the post-FMT adult group and adult pooled sample. However, after these mice had been inoculated with the adult pooled sample, they clustered with the aged post-FMT group. The differences in the faecal microbiotas between the adult and aged mice post-FMT were not as pronounced as the differences seen between the adult and aged groups in our initial study; as such a less stringent Benjamini–Hochberg cut-off (P<0.1) was used when analysing these data (Fig. 1). This less-pronounced change is unsurprising as the adult and aged mice in the initial study had not been subject to any intervention, however it is becoming clear that even subtle changes in the gut microbiome are associated with phenotypic changes in the host [34] and the same may be true of cognitive traits. Only four genera (*Prevotellaceae, Faecalibaculum, Lachnospiraceae* and *Ruminococcaceae*) were found to be significantly differentially abundant in the faeces of the post-FMT adult and post-FMT aged animals (Fig. 2c). Few significant associations were seen between the faecal microbiomic and metabolomic data (Fig. 2d).

**Figure 2.**
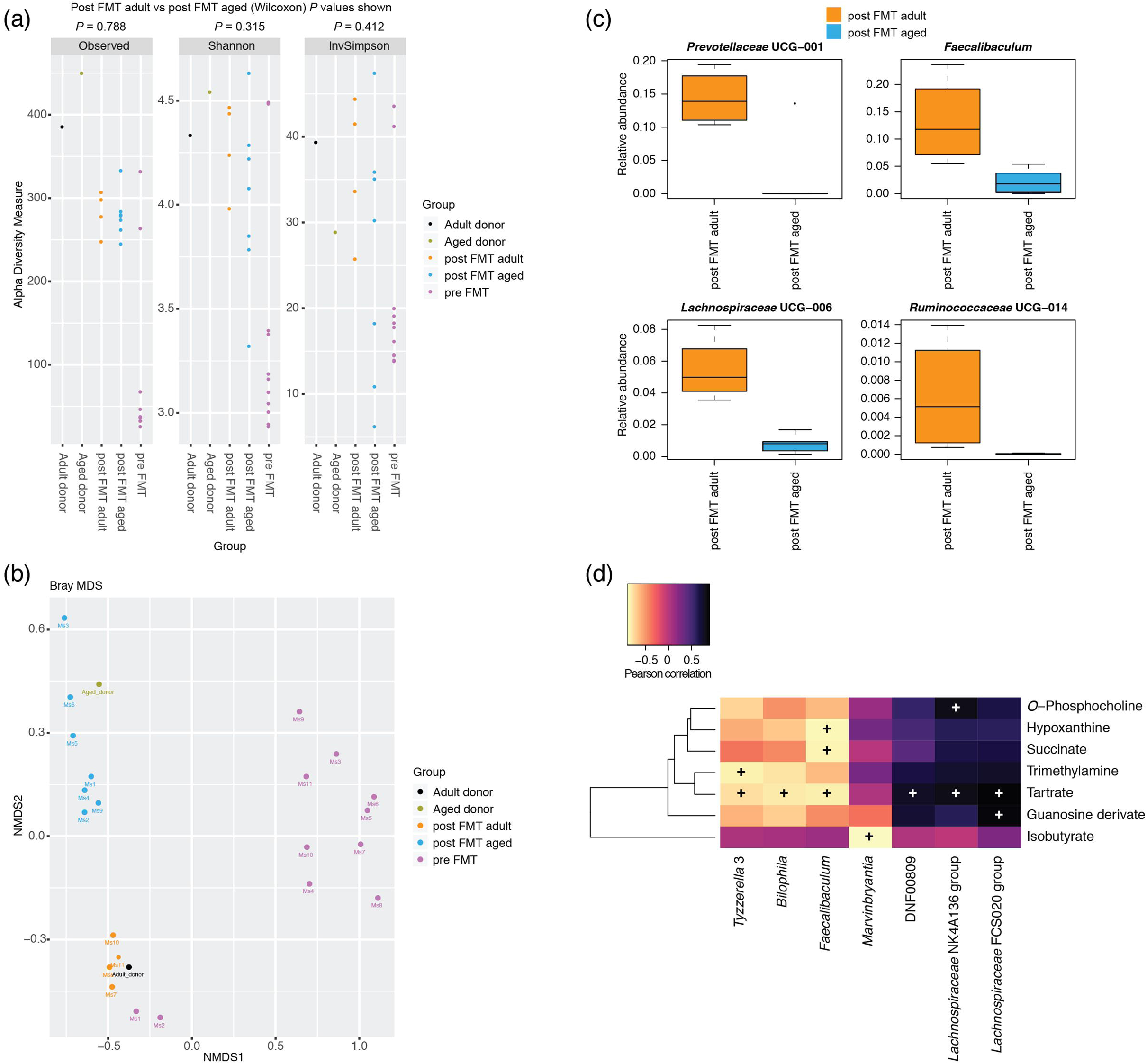
Effect of FMT on the faecal microbiotas of adult mice. (a) Measures of alpha diversity among the pooled adult (n=1) and aged (n=1) samples, the adult mice pre FMT (n=11) and the adult mice after FMT with adult (n=4) and aged (n=7) faeces. (b) MDS plot of a Bray Curtis assessment of beta diversity. Data presented are for ASVs present in more than two animals and prevalence threshold of 1 % at the genus level. (c) Box plots for the genera that were significantly different (Welch’s t test and Wilcoxon; P<0.1, Benjamini– Hochberg) between post FMT adult (n=4) and post FMT aged (n=7) mice based on ALDEx2 analyses. (d) Pearson correlation of faecal microbiomic and metabolomic data. Only rows/columns that contained significant data (P<0.1, Benjamini–Hochberg) are shown.

### FMT from aged mice to adult recipients results in decreased spatial learning and memory but does not affect motor activity or anxiety-like behaviour

In the first set of experiments, we investigated the impact of FMT from aged mice to adult recipients to a series of spatial learning and memory tests. First, the Barnes maze test was used to assess memory and spatial learning. Following the training trials, a retention test was conducted in which the escape tunnel was removed and the latency before moving to the position of the former escape tunnel for the first time was measured. We observed that the average primary latency was significantly higher for FMT-adult recipients of microbiota from aged donors (FMT-aged) than the other control groups either left untreated or colonized with microbiota from adult, age-matched donors (FMT-adult) (41.7 ± 3.5 s; 30.5 ± 3.9 s and 23.5 ± 6.7 s, respectively; P<0.05) (Fig. 3a). Yet, during the retention test, FMT-aged mice spent less time in the target quadrant (28.9 ± 2.6 s in comparison to control 46.5 ± 6.3 s, and FMT-adult 51.8 ± 10.2 s; P<0.05) that contained the escape tunnel compared to control group (Fig. 3b; heat map in Fig. 3c) confirming a direct impact of the gut microbiota on memory and spatial learning. Learning and memory was further evaluated by the novel object recognition test. Significant differences between groups were observed in the time spent exploring two different objects (Fig. 3d). In particular, the time spent exploring novel and familiar objects was monitored; the object recognition was assessed by a defined discrimination index. Control groups, either untreated or FMT-adult mice, preferred the novel object more than the familiar one, whereas FMT-aged mice showed a significant reduction in the time spent exploring the novel object compared to control (0.23 ± 0.04 and 0.46 ± 0.05, respectively; P<0.01) suggesting reduced discrimination as a consequence of impaired memory capabilities. This is further evidenced by the heat maps (Fig. 3e).

**Figure 3.**
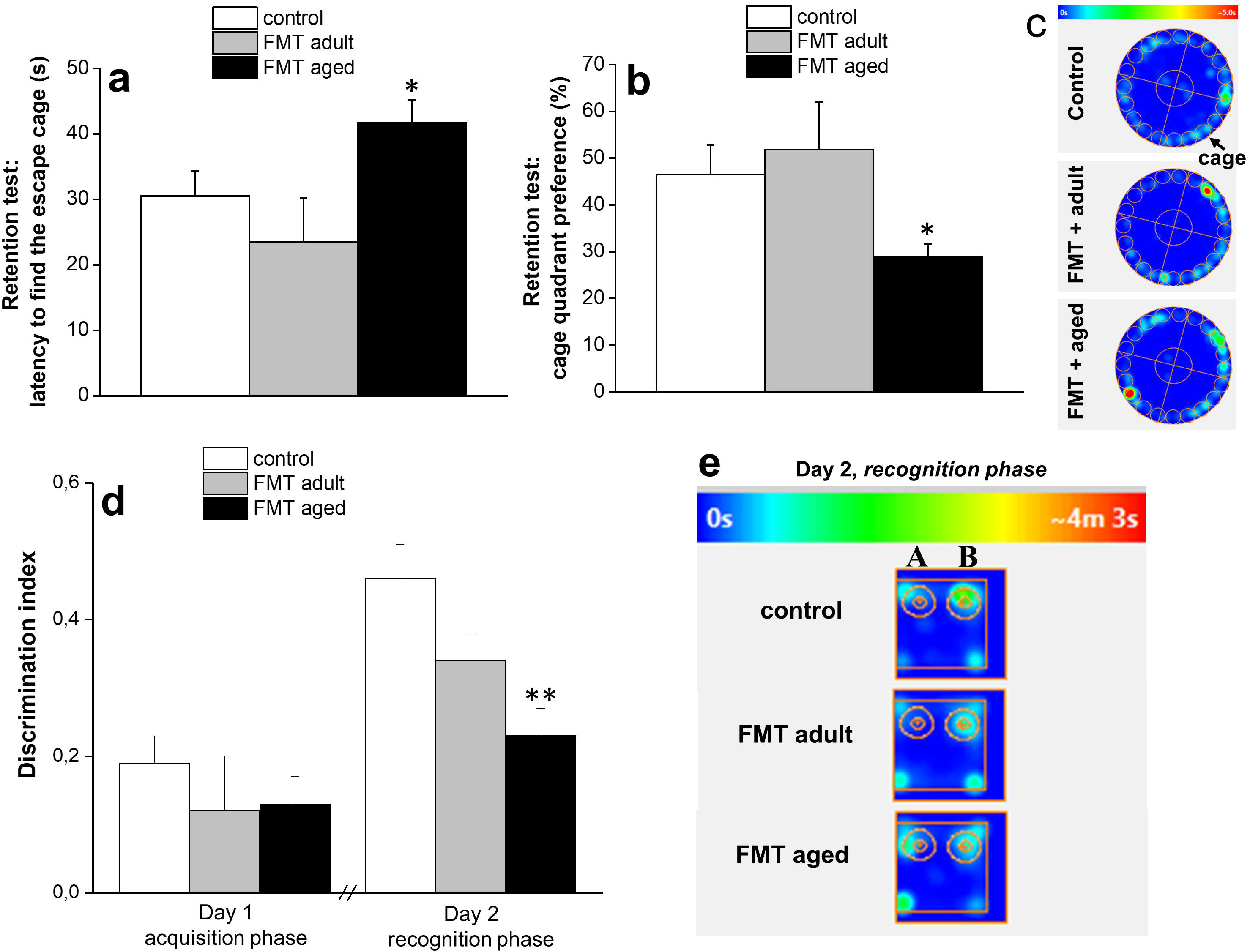
Effect of FMT from adult and old mice on spatial learning and memory. Barnes Maze test (Fig 3a-c): mice were trained to find the cage for 4 consecutive days (twice daily; 2 trials). The average primary latency (Fig 3a) was significantly higher for adult recipients of microbiota from aged donors (FMT-aged) than the other control groups either left untreated or colonized with microbiota from adult, age-matched donors (FMT-adult). Furthermore, during the retention test, FMT-aged mice spent less time in the target quadrant that contained the escape tunnel compared to control groups (Fig. 3b; heat map in Fig. 3c). The values represent the mean ± SEM for each group (n=10-12 mice/group). *P<0.05 vs control animals and FMT-adult. The novel object recognition test (Fig. 3d-e): on day 1, mice were exposed to two similar objects (A+A); on day 2, animals were re-exposed to the testing area containing one novel object (A+B). The time spent by the animals exploring each object was recorded. The discrimination index, calculated as (TB-TA)/(TB+TA), was used to assess the preference for the novel object. Control groups, either untreated or FMT-adult mice, preferred the novel object more than the familiar one, whereas FMT-aged mice showed a significant reduction in the time spent exploring the novel object (heat map in 3e) suggesting reduced discrimination as a consequence of impaired memory capabilities. The values represent the mean ± SEM for each group (n=10-12 mice/group). **P<0.01 vs control animals.

Given that the microbiota has been reported to affect locomotor activity and anxiety-like behaviour in animal models [5, 35], we then assessed whether FMT from aged animals to adult recipients could also affect these behavioural manifestations. For this purpose, we employed two validated tests: the open field and the elevated plus maze. In the open field task, no significant difference in the distance travelled by both FMT-aged and FMT-adult was observed when compared to the untreated control group indicating no significant motor impairments of the animals (Fig. 4a). However, it is worth noting that FMT-aged mice displayed a tendency to prefer the periphery or the corners of the arena instead of the centre (Fig. 4b). The representative tracks of movement patterns are depicted in Fig. 4c-e (ANY-maze software). In the elevated plus maze FMT-aged mice did not display significant differences in time spent in either arms of the maze compared to control groups (Fig. 4f-g) with most of the animals spending the majority of the time in the closed arms irrespective of the FMT treatment.

**Figure 4.**
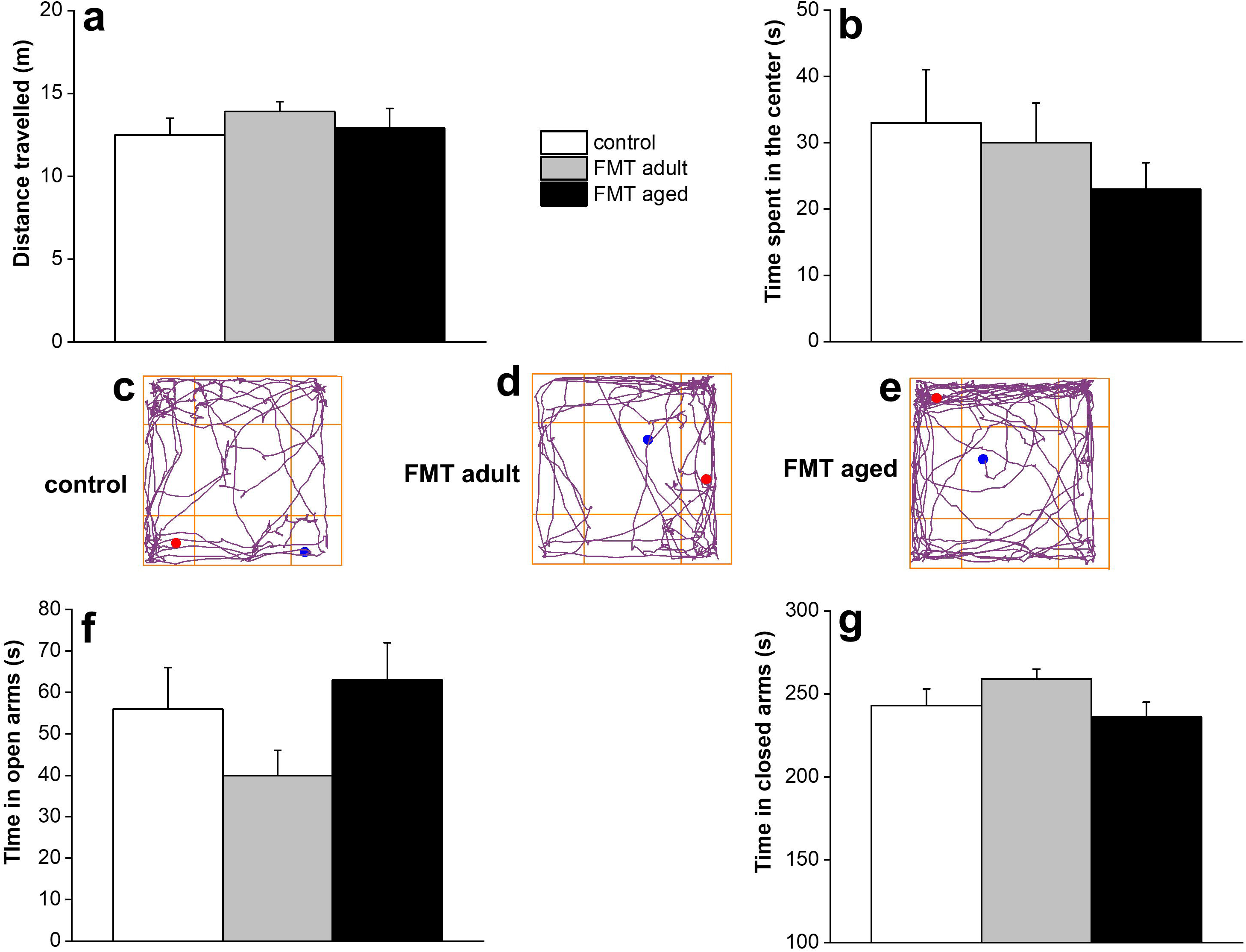
Effect of FMT from adult and old mice on locomotor and explorative activity and anxiety-related behavior. Open Field Test did not show significant difference in the distance travelled by both FMT-aged and FMT-adult was observed when compared to the untreated control group indicating no significant motor impairments of the animals (Fig. 4a). However, FMT-aged mice displayed a tendency to prefer the periphery or the corners of the arena instead of the center (Fig. 4b). The representative tracks of movement patterns are depicted in Fig. 4c-e (ANY-maze software Furthermore, in the elevated plus maze, FMT-aged mice did not display significant differences in time spent in either arms of the maze compared to control groups (Fig. 4f-g). The values represent the mean ± SEM for each group (n=10-12 mice/group).

### FMT from aged donors alters protein expression in the hippocampus of adult recipients

The observation that FMT from aged donors to adult recipients could alter behavioural patterns of transplanted mice prompted us to investigate the molecular mechanisms underlying such changes. To this extent, a one-shot label-free quantitative proteomics approach was employed, targeting the hippocampus, a brain region known to play an important role in learning and memory and to undergo several changes with the advancing of age [36]. The analyses resulted in the quantitation of 2180 protein groups and 16083 unique peptides (Supplementary Table S1). The Pearson correlation coefficients between samples and technical replicates were 0.99, indicating a high reproducibility and confidence of the resulting data (Supplementary Fig. S3A). The volcano plot was obtained by two-sided t test of the two groups: FMT aged faeces into adult mice related to the adult FMT into adult mice (Supplementary Fig. S3B). 140 proteins were differentially regulated, 52 overexpressed and 88 down-expressed, of which 47 proteins were differentially regulated with a fold change higher than 1.5 (Table 1, Supplementary Table S1). Furthermore, western blot analyses of two selected proteins implicated in the behavioural phenotype of transplanted mice, the microtubule-associated protein MAPT (FMT-aged versus FMT-adult ratio: 2.38) and tRNA-splicing ligase RtcB homolog Rtcb (FMT-aged versus FMT-adult of 0.46) confirmed the patterns identified by the label-free proteomics approach (Supplementary Fig. S4 and Supplementary Fig. S5).

**Table 1.**
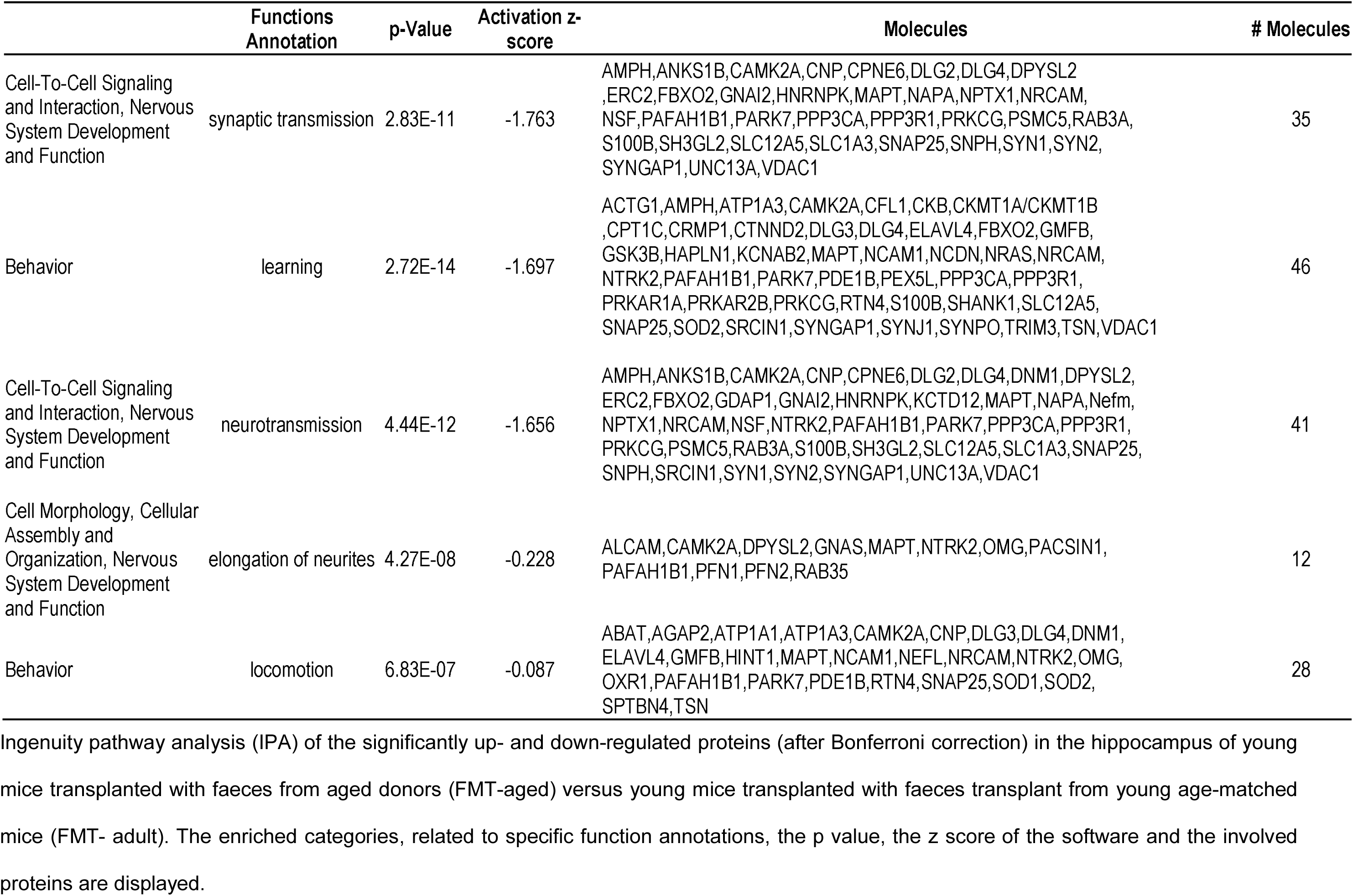
Ingenuity Pathway Analysis (IPA)

Subsequent protein network analyses highlighted several differentially expressed proteins in the hippocampus of adult mice that received microbiota from aged donors compared to adult mice that received microbiota from age-matched adult donors. Table 1 summarises the main enriched categories. Cellular signalling during nervous-system development and related to synaptic transmission was described by 35 molecules, differentially expressed, indicating a downregulation during the transplant (z score −1.7). Neurotransmission was also found downregulated by 41 differentially expressed molecules. Remarkably the changes in the expression of a total of 87 learning-related proteins and cognition tests in FMT-aged mice pointed to a decline of brain functions as seen during the physiological ageing process (Fig. 5). As a corollary, the extent of the effects of the shifts of microbiota on health and disease in ageing is further stressed by the observation that a large set of proteins involved in lipid metabolism were down-regulated (Supplementary Fig. S6). This latter evidence suggested that shifts of microbiota might underpin the detrimental effects of ageing on multiple health-critical functions. Thus, the single label-free run approach resulted in an in-depth hippocampus protein characterization and above all in the identification of differentially regulated proteins involved in key pathways of ageing-related processes.

**Figure 5.**
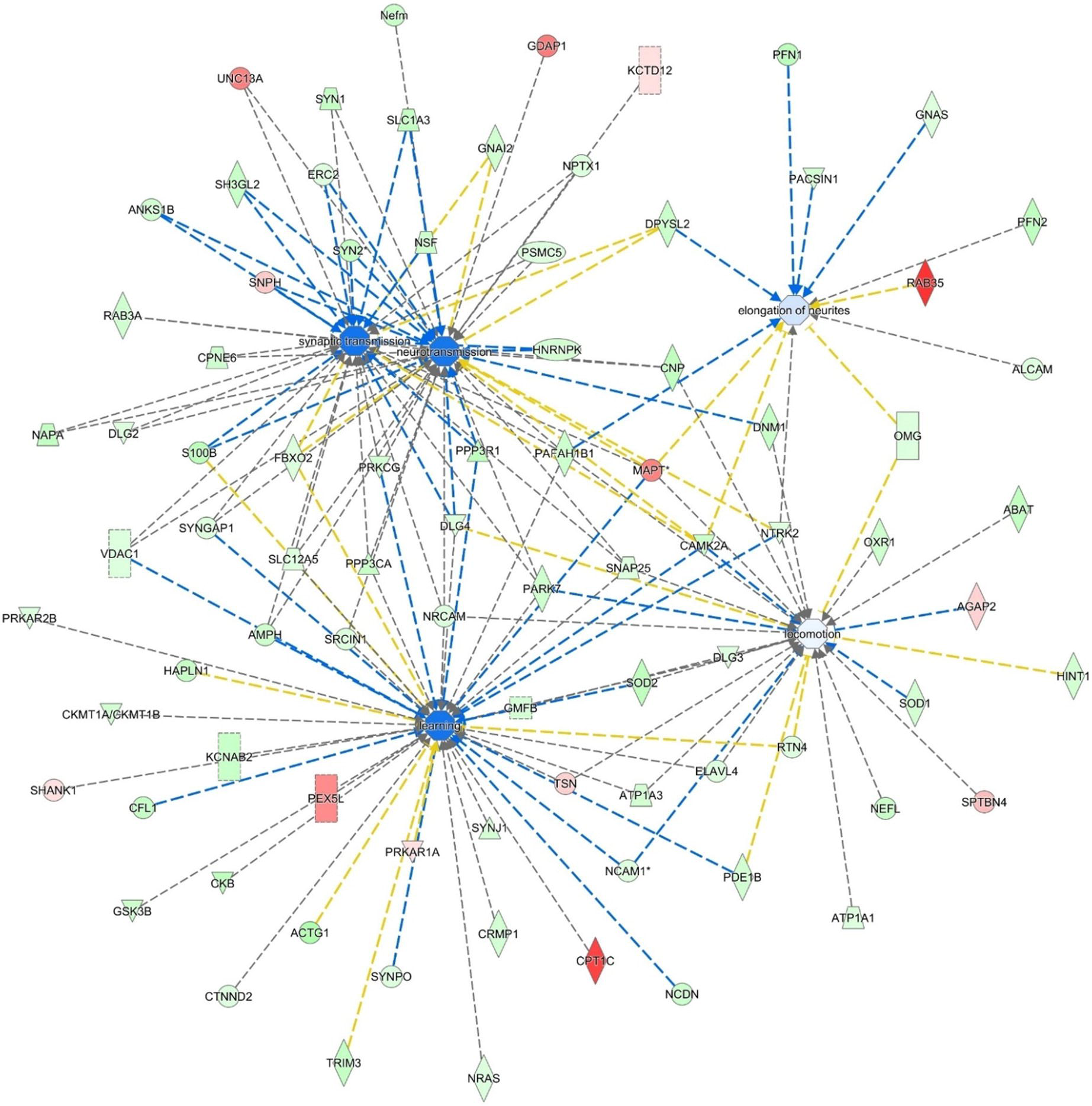
Ingenuity Pathway Analysis (IPA). IPA of the significantly up- and down-regulated proteins (after Bonferroni correction) for faeces from aged donors transplanted in adult mice versus faeces from adult donors transplanted in adult age-matched mice, in hippocampus tissue. The circles represent the main network node and the blue colour the significantly downregulated nodes. The up-regulated proteins are marked red, while those that that were down-regulated are marked in green. The intensity of the colour relates to fold-change (light to dark colour = small to large fold change). The symbols shown in the network are explained at http://www.qiagenbioinformatics.com/products/ingenuity-pathway-analysis.

### FMT from aged donors into adult recipients triggers phenotypic changes in glia cells of the hippocampus fimbria but did not affect gut permeability or levels of circulating cytokines

It has been suggested that FMT from aged donors triggers systemic inflammageing in adult recipients [37]. Consequently, we investigated whether FMT from aged donors triggered increased systemic levels of pro-inflammatory cytokines and intestinal permeability in adult recipients. We report that neither gut permeability nor plasma levels of inflammatory cytokines (Supplementary Fig. S7A and B) changed following FMT. Also, FMT did not induce increase in glial fibrillary acidic protein (GFAP) expression in astrocytes of hippocampus regions (Fig. 6A-H) suggesting the lack of an overt neuroinflammatory response. On the other hand, a significant (P = 0.0168) increase in the expression of F4/80, a typical trait of the ageing brain [38], was observed in glia cells in the white matter of the hippocampus fimbria (Fig. 6J-L).

**Figure 6.**
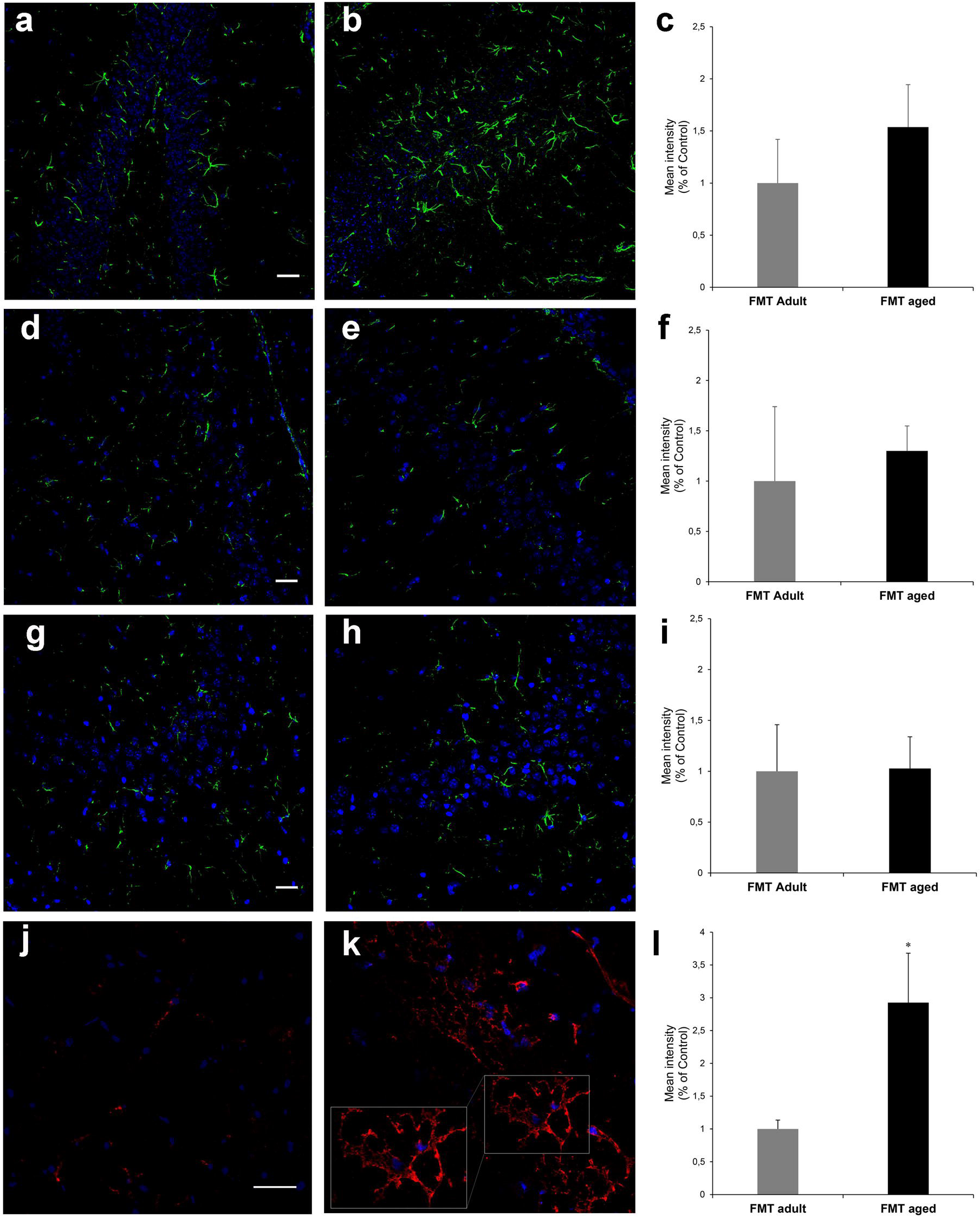
Post-FMT levels of GFAP and F4/80 in hippocampal areas. Representative images acquired at the confocal microscope with anti-GFAP antibody (green) in the hippocampal regions (a-b, d-e, g-h) and anti-F4/80 antibody (red) in the fimbria (j-k) of FMT adult mice (a, d, g, j) or FMT aged mice (b, e, h, k). Nuclei have been counterstained with ToPro-3 (in blue). No difference was observed in the expression of GFAP in the dentate gyrus (a-c), CA4 (d-f) and CA3 (g-i). Bars in c-f and I represent fluorescence intensity (mean ± SEM). On the contrary, glia cells of the hippocampus fimbria showed an increased expression of F4/80 (j-k), the latter being a feature of the ageing brain. Bars in (l) show the levels of fluorescence intensity (mean ± SEM) in FMT-adult and FMT-aged recipients (asterisk indicates p=0.0168) (bars 30 μm) (N=3 mice/group)

## Discussion

The decline of cognitive and behavioural functions in ageing is one of the most detrimental effects of ageing [34]. In particular, a deficit of hippocampal-dependent functions, including spatial learning and memory is a distinctive trait of ageing [17]. Although the role of the gastrointestinal tract and gut microbiome on the development and function of the CNS has become evident in the past few years [2-6], the extent of the impact of the age-associated shits of the gut microbiota on specific functions of the CNS remained to be determined. Here we report that FMT from aged donors affects memory and spatial learning in transplanted adult mice, a phenomenon led through a differential expression of proteins involved in maintenance of synaptic plasticity and neurotransmission in the hippocampus area. On the other hand, anxiety, explorative prowess and locomotor activity were not affected by the FMT procedure.

Although the presence of specific microbial families and genera have been associated with cognitive decline, anxiety behaviours and affective disorders [39], to our knowledge this is the first evidence showing that FMT significantly affects important cognitive functions of the CNS in ageing. Such observations prompted us to investigate the potential underlying mechanisms and whether the observed changes in cognition and behaviour may be triggered through the modification of proteins relevant for CNS function.

A label-free quantitative proteomics approach showed a significant modifications of protein expression in the hippocampus area. In particular, the variation of the expression of proteins, such as D-dopachrome decarboxylase (D-DT), neuronal membrane glycoprotein M6-a (Gmp6A) and M6-b and Ras/Rap GTPase-activating protein SynGAP was of significance. Indeed, alterations in the expression of these proteins in the CNS have been implicated in important CNS functions and disturbances, ranging from filopodium formation, synaptogenesis, learning disability, behavioural anomalies and neuronal plasticity to neurodegenerative disorders [40]. In addition to these observations, it has been suggested that changes in the microbiota are associated with increased gut permeability and systemic inflammation that may play an important role in age-associated alteration of cognitive and behaviours in ageing [41]. With regard to inflammation it has been described that FMT from aged donors to young germ-free recipients triggered systemic inflammageing including enhanced CD4^+^ T cell differentiation and distribution of several Th1 and Treg subsets, in particular in the systemic compartment [37]. However, the latter report fell short of measuring circulating pro-inflammatory cytokines and cognitive/behavioural functions in the recipients. In our experiments, a small non-significant increase was observed for IL-1β and TNF-α. Consistent with the lack of a significant inflammatory response is our observation that levels of GFAP did not increase in various areas of the hippocampus. However, a direct effect of pro-inflammatory cytokines on the protein expression and function of the hippocampus cannot be ruled out. It is plausible that a drastic increase of systemic levels of cytokines is not required and that only a very minor increase of specific systemic cytokines may suffice to trigger the changes observed here. The latter hypothesis is supported by the phenotypic change of the microglia, resident immunocompetent cells in the CNS that respond to a plethora of inflammatory stimuli [42]. Indeed, adult recipients that had received FMT from aged donors showed a significant increase in the expression of the marker F4/80, a specific trait of the ageing brain [38]. The latter result is of potential importance. Indeed, microglia cells monitor and regulate specific neuronal elements including synapses [43] and neuronal cell bodies [44] and in so doing they play an active role in neuronal surveillance in homeostatic conditions as well as in response to disease and injury. Currently, whether the FMT-induced phenotypic change of the microglia is paralleled by a reduced capability to contribute to be determined.

Furthermore, integration of faecal microbiomic and metabolomic data showed a clear association between lower levels of SCFAs and decreased representation of obligate anaerobes such as the *Lachnospiraceae* and *Ruminococcaceae* that have been associated with production of SCFAs by the human faecal microbiota (Supplementary Fig. S1 and Supplementary Fig. S2). It should, however, be noted the biological relevance of increased levels of faecal propionate, butyrate and isobutyrate being significantly correlated with increased relative abundance of lactic acid bacteria such as *Staphylococcus, Jeotgalicoccus* and *Facklamia* is currently unknown (Supplementary Fig. S1 and Supplementary Fig. S2), as our knowledge as to the ability of these species to produce SCFAs other than acetate or lactate from fermentation of carbohydrate or protein sources is poor as is our knowledge on cross-feeding potential of the mouse gut microbiota.

In addition to these aforementioned genera, a strong decrease in *Prevotellaceae* was also observed following FMT from aged donors in adult recipients. We previously reported higher levels of *Prevotellaceae* and *Ruminococcaceae* in *APOE3/E3* and *APOE2/E3* genotype carriers, respectively, relative to *APOE4* carriers [26], one of the strongest prevalent risk factors for neuropathology and Alzheimer’s disease [45]. Whilst the under-representation of *Prevotellaceae* has been previously reported to diminish the levels of health-promoting neuroactive SCFAs in humans [46] and the biosynthesis of thiamine and folate, two vitamins decreased in Parkinson’s disease [47, 48], depletion of *Ruminococcaceae* has been associated with Alzheimer’s disease [49]. Interestingly, a previous study showed that SCFAs may regulate host serotonin biosynthesis [50], a multifaceted neurotransmitter modulating cognition, learning and memory, along with numerous physiological processes [51]. Finally, the role of the vagus nerve on microbiota-mediated alterations of the hippocampus area cannot be rule out. This cranial nerve exerts an important role in gut-brain communication [52] and it has been shown that it mediates the effects of orally delivered probiotic strains on aspects of behaviour [53]. Future experiments of selective vagotomy will help to elucidate the role of the vagus nerve in the microbiota-mediated decline of spatial and learning memory.

Overall, these results further highlight the paramount importance of events taking place in the GI-tract in health and disease. In particular, they point to a direct impact of age-associated shifts of the intestinal microbiota composition on the decline of key functions of the CNS. This notion, along with recently collected evidence that correcting age-associated shift of microbiota profile is beneficial for health and life expectancy [54], lends support to the hypothesis that microbe-based approaches that aim to restore a young-like microbiota might improve cognitive function and in so doing the quality of life of the elderly, an ever-increasing demographic segment of modern societies.

## Acknowledgments

The authors wish to thank S. Deakin, UEA Norwich for his technical help. This work was partly supported by the research grants from the Ente Cassa di Risparmio di Firenze (to CN), core grants from the University of Florence, Florence, Italy (to CN and MG) and by the Biotechnology and Biological Sciences Research Council (BBSRC), UK, Institute Strategic Programme Grant (BB/J004529/1) (to AN). This work used the computing resources of the UK MEDical BIOinformatics partnership (UK Med-Bio), which was supported by the Medical Research Council (MR/L01632X/1 to LH). This manuscript has been released as a pre-Print at bioRxiv (doi:https://doi.org/10.1101/866459)

## AUTHOR CONTRIBUTIONS

CN, AdA, AN, DV, LM conceptualized and designed the experiments and analytical approaches; LH processed, analysed and interpreted 16S rRNA gene sequence data. GLG processed, analysed and interpreted metabolomics data; PG, CG designed cognition and behaviour experiments; EL, ALM, JVB, EB, MR, AP, GL, AA carried out experiments and analysed data. CN, DV, AdA, LM, MG and LH wrote the paper with contributions from all authors. All authors read and approved the final manuscript.

## Figure legends

**Supplementary Figure S1. Pearson correlation of faecal microbiomic and metabolomic data for adult and aged mice**. ALDEx2 was used to correlate the datasets.+, P<0.05 (Benjamini-Hochberg). Only rows/columns containing significant data are shown: all results are shown in Supplementary Fig 2 (8 mice/group).

**Supplementary Figure S2. Pearson correlation of faecal microbiomic and metabolomic data for adult and aged mice**. ALDEx2 was used to correlate the datasets. +, P<0.05 (Benjamini-Hochberg). All data are shown: significant results were used to generate Supplementary Figure 1 (8 mice/group).

**Supplementary Figure S3. Post-FMT quantitative analysis of proteins in the hippocampus**. Volcano plot of quantified proteins in hippocampus tissue (A). Differentially regulated proteins due to faeces from aged mice transplanted into adult mice (T) versus faeces from adult mice transplanted into adult age-matched mice (C) are showed (T/C). The proteins in red are up regulated and in green down regulated. Scatter Plot of protein intensities (B) obtained by label free quantitation by MaxLFQ in MAxQuant, showing the Person correlation coefficients between biological and technical replicates of analysed samples.

**Supplementary Figure S4. Western Blot analysis for Mapt in the hippocampus of FMT-treated mice**. Polyacrylamide (12%) gel stained with blue Coomassie with a representative image of molecular weight marker with relevant kDa (a). GAPDH visualized bands and merged with nitrocellulose membrane (b). Mapt visualized bands and merged with nitrocellulose membrane (c). In (d) a representative histogram shows levels of analysed protein both in FMT-Y and FMT-Atreated and control animals. Lane 1 (positive control, SH-SY5Y cell line); lane 2 (negative control, H292 cell line); lane 3 (aged mouse hippocampal proteins); lane 4 (adult mouse hippocampal protein); lane 5 (FMT-aged hippocampal proteins); lane 6 (FMT-adult hippocampal proteins). Mapt protein was detected approximately at 50 kDa (right blots). GAPDH (37 kDa) was used as housekeeping

**Supplementary Figure S5. Western Blot analysis for Rtcb in the hippocampus of FMT-treated mice**. Polyacrylamide (12%) gel stained with blue Coomassie with a representative image of molecular weight marker with relevant kDa (a). GAPDH visualized bands (panel b, left) and merged with nitrocellulose membrane (panel b, right). Rtcb visualized bands (panel c, left) and merged with nitrocellulose membrane (panel c, right). In (d) representative histogram shows levels of analysed protein both in FMT-Y and FMT-A treated and control animals. Lane 1 (positive control, SH-SY5Y cell line); lane 2 (negative control, mouse adipose tissue); lane 3 (aged mouse hippocampal proteins); lane 4 (adult mouse hippocampal protein); lane 5 (MT-aged hippocampal proteins); lane 6 (MT-adult hippocampal proteins). Rtcb protein was detected approximately at 56 kDa (right blots). GAPDH (37 kDa) was used as housekeeping (left panel). Molecular weight (mw) used was SHARPMASS VII.

**Supplementary Figure S6. Ingenuity pathway analysis (IPA)**. IPA analysis of the significantly up- and down-regulated proteins (after Bonferroni analysis) for faeces from aged donors transplanted in adult mice versus faeces from adult donors transplanted into adult age-matched mice, in hippocampus tissue. The circles represent the main network node and the blue colour the significantly down regulated. The up-regulated proteins are marked in red, while those that that were down-regulated are marked in green.

**Supplementary Figure S7. FMT from aged donors does not affect either gut permeability or circulating cytokines**. Mice were orally administered with a solution of FITC-dextran and plasma levels measured after 45 minutes. No differences were observed between FMT-adult and FMT-aged recipients. Plasma samples were also used to evaluate levels of circulating cytokines; also in this case we failed to observe any significant change of circulating pro- and anti-inflammatory cytokines ion both group of FMT-treated mice (n=8 mice/group).

