## Supplementary figures and images for "Faecal microbiota transplant from aged donor mice affects spatial learning and memory via modulating hippocampal synaptic plasticity- and neurotransmission-related proteins in young recipients"

### Supplementary Figure S1.JPG

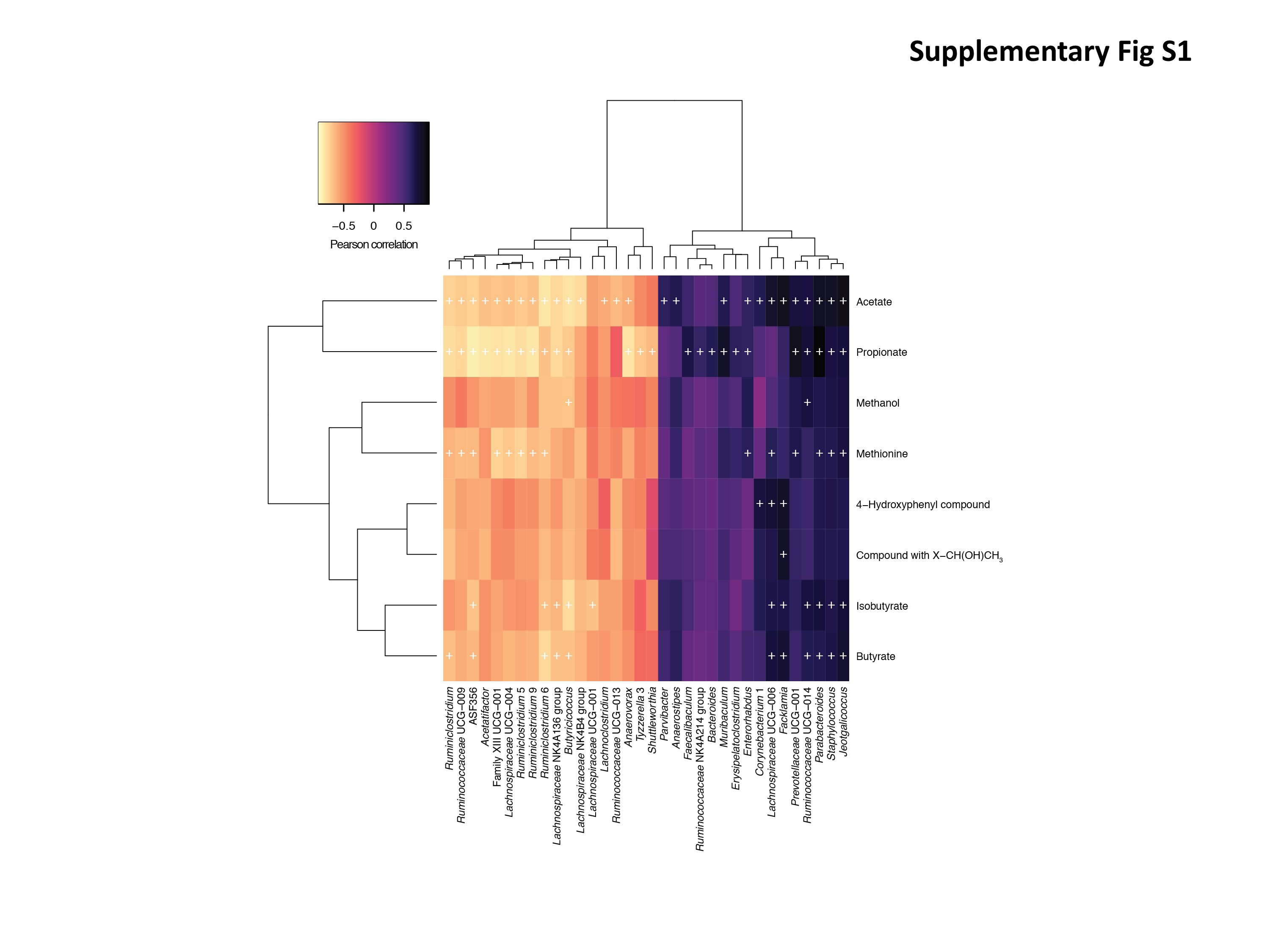

### Supplementary Figure S2.JPG

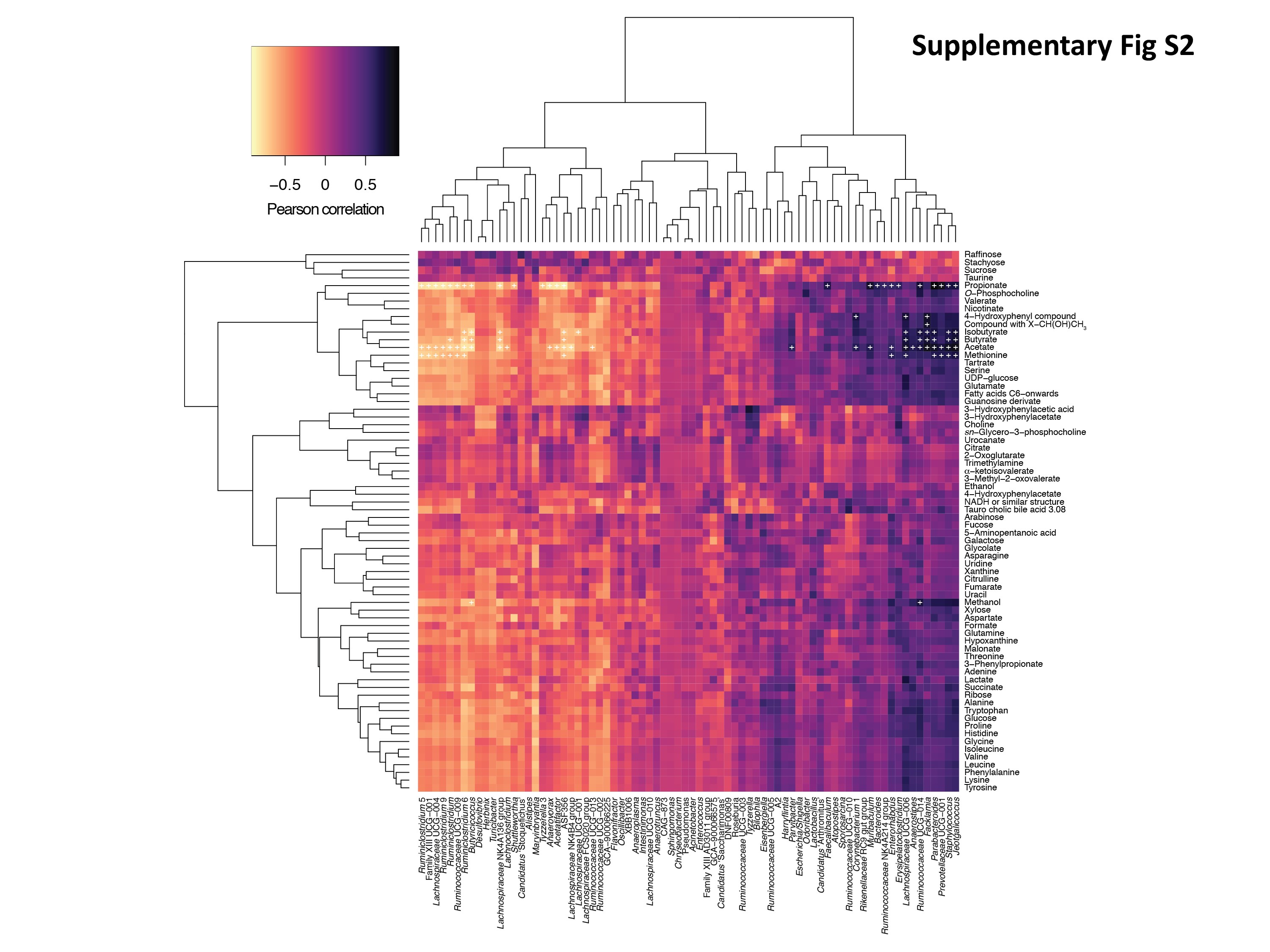

### Supplementary Figure S3.JPG

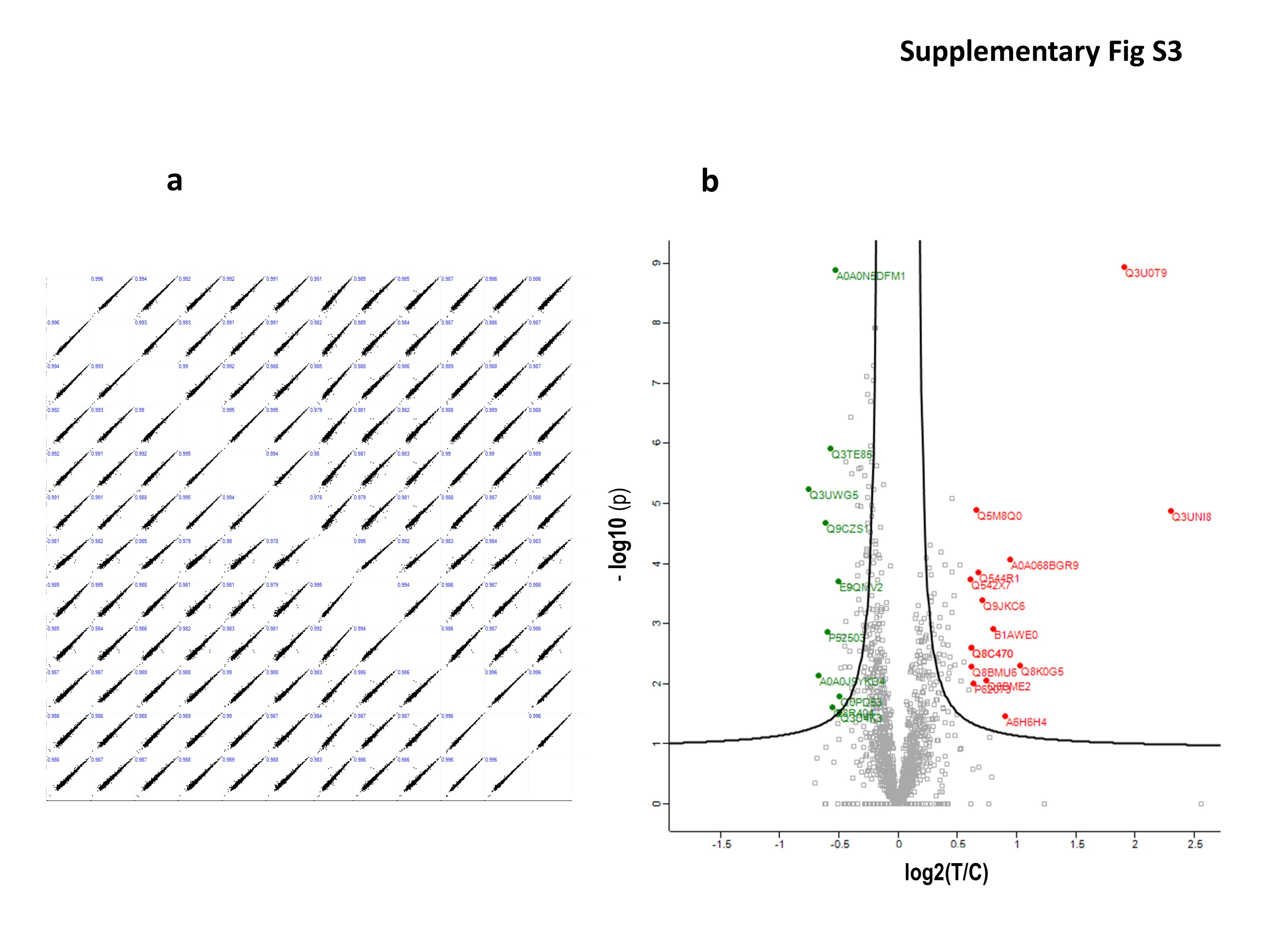

### Supplementary Figure S4.JPG

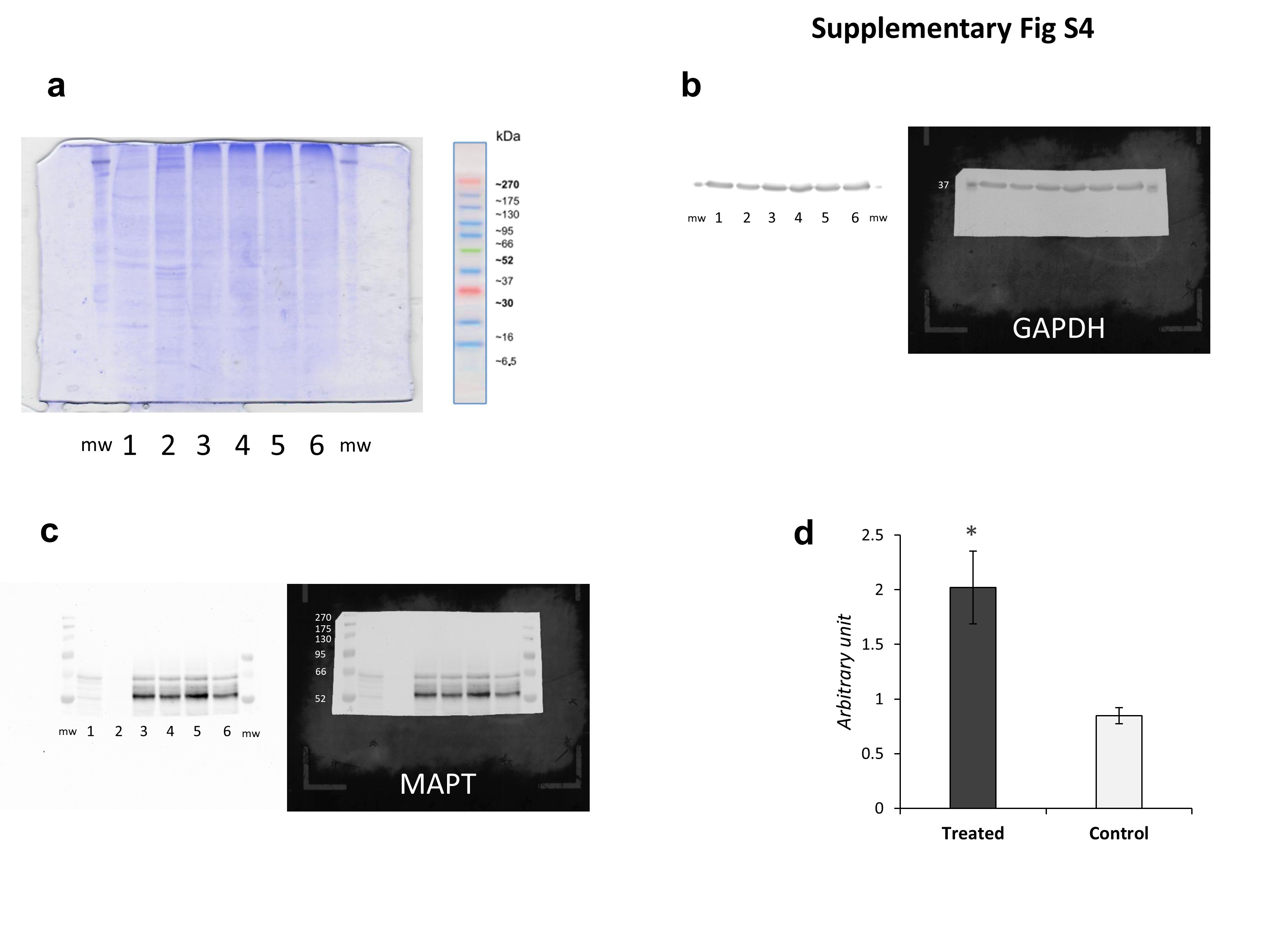

### Supplementary Figure S5.JPG

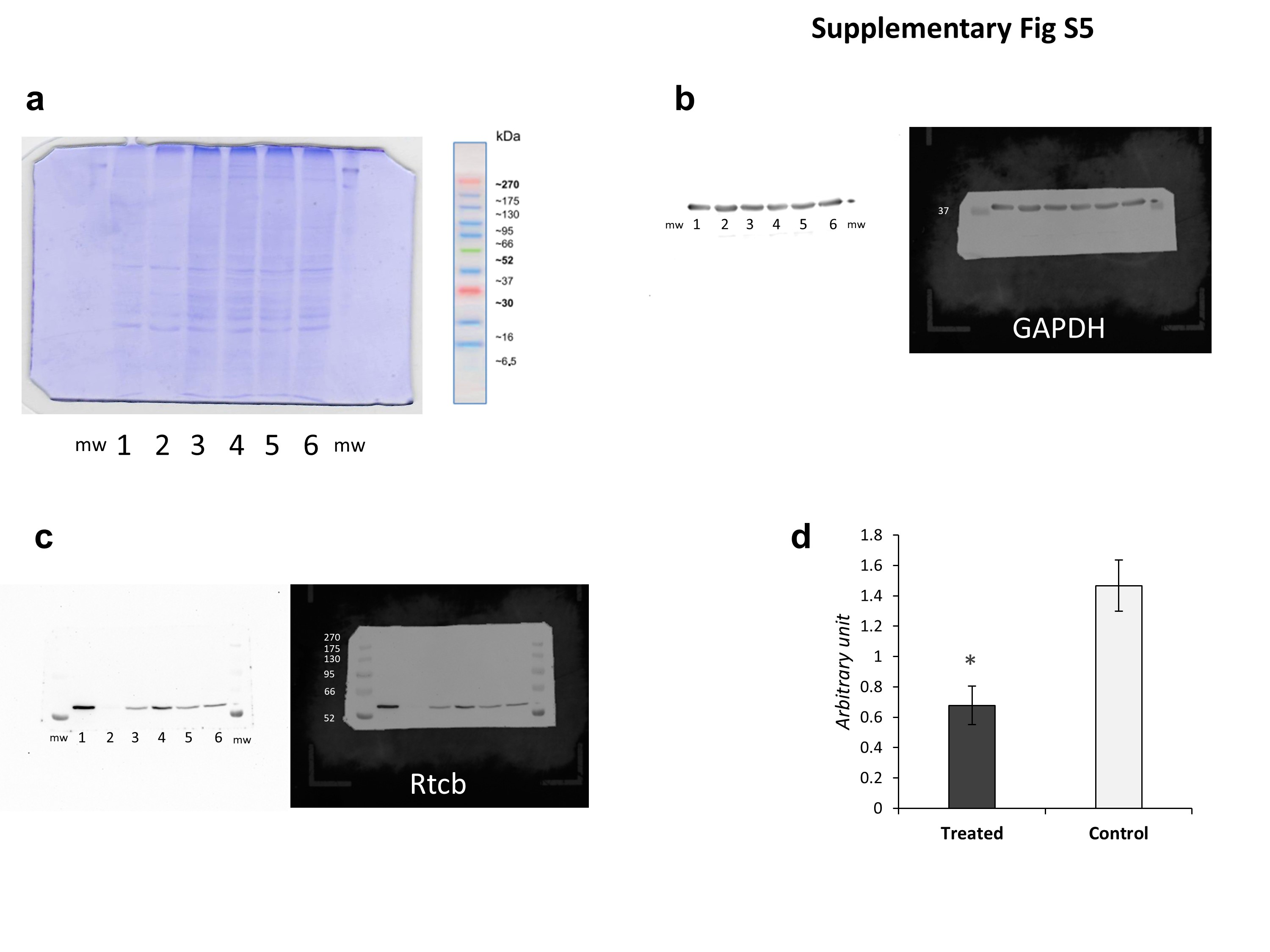

### Supplementary Figure S6.JPG

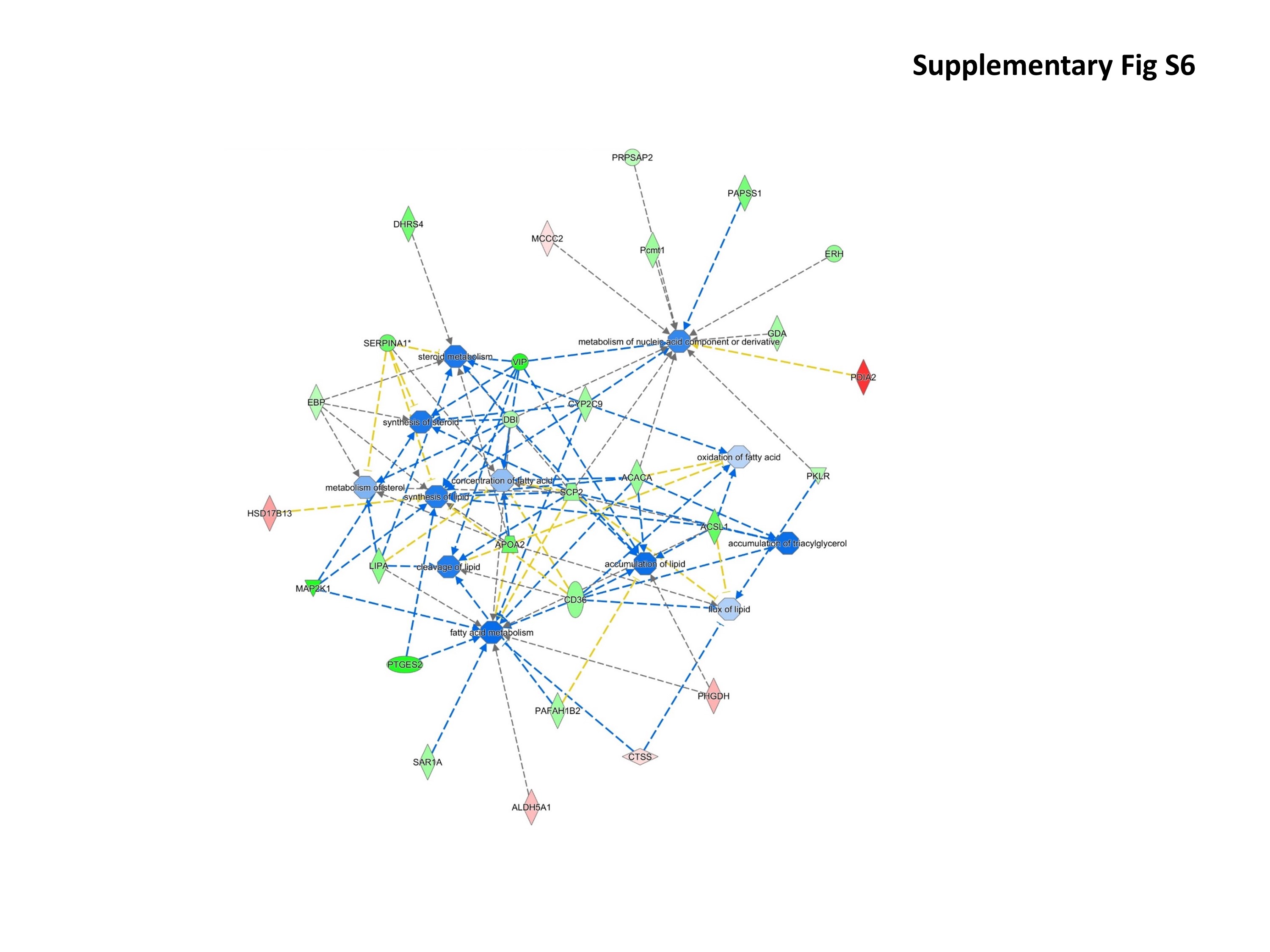

### Supplementary Figure S7.JPG

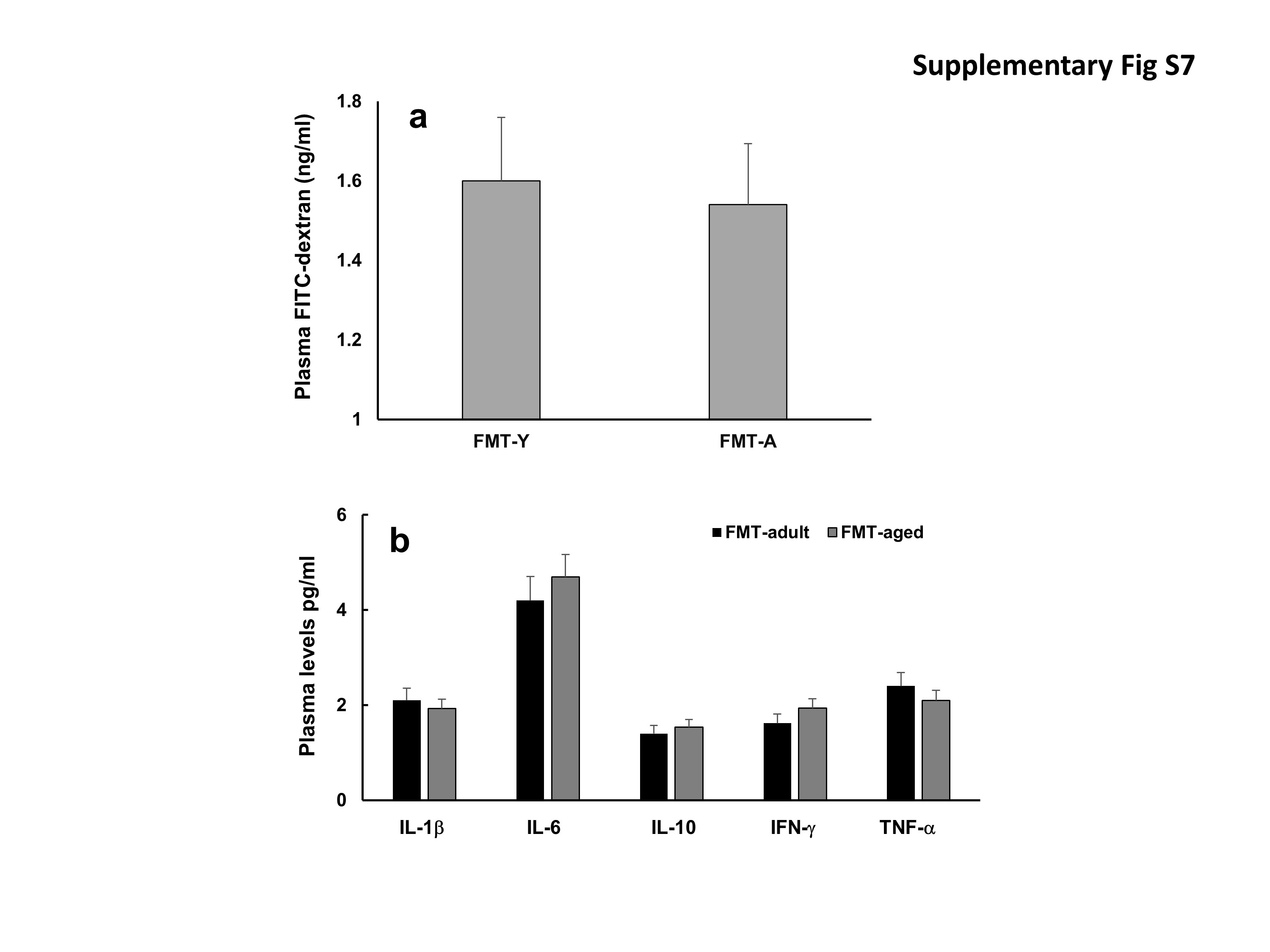
